# Reduced brain glutathione levels during normal aging are associated with visuospatial memory

**DOI:** 10.1101/2023.08.13.553100

**Authors:** Xin Hu, Keyu Pan, Min Zhao, Jiali Lv, Jing Wang, Xiaofeng Zhang, Yuxi Liu, Yulu Song, Aaron T. Gudmundson, Richard A.E. Edden, Fuxin Ren, Tao Zhang, Fei Gao

## Abstract

During aging, the brain is subject to greater oxidative stress (OS), which is thought to play a critical role in cognitive impairment. Glutathione (GSH), as a major antioxidant in the brain, can be used to combatting OS. However, how brain GSH levels vary with age and their associations with cognitive function remain unclear. In this study, we combined point-resolved spectroscopy and edited spectroscopy sequences to investigate GSH levels in the anterior cingulate cortex (ACC), posterior cingulate cortex (PCC), and occipital cortex (OC) of 276 healthy participants (166 females, age range 20–70 years) and examined their relationships with age and cognitive function. The results revealed decreased GSH levels with age in the PCC among all participants. Notably, the timecourse of GSH level changes in the PCC and ACC differed between males and females. Additionally, positive correlations were observed between GSH levels in the PCC and OC and visuospatial memory. Taken together, these findings enhance our understanding of the brain GSH timecourse during normal aging and associations with sex and memory, which is an essential first step for understanding the neurochemical underpinnings of OS-related diseases.

## Introduction

Glutathione (GSH) is a tripeptide consisting of glycine, glutamate, and cysteine present in almost all cells. In addition to having roles in numerous physiological functions, GSH is an important antioxidant for antioxidant defense ^1^. Several studies have suggested that low levels of reactive oxygen species (ROS) contribute to maintaining normal cellular function. However, accumulation of ROS causes oxidative stress (OS), resulting in lipid peroxidation, protein oxidation, and DNA damage that ultimately impair cellular function ^2^. The brain is particularly susceptible to OS, as it consumes 20% of the body’s oxygen despite accounting for only 2% of total body mass ^3^. Despite awareness of the crucial role of GSH, how GSH levels in the brain change with age remains unclear.

The majority of rodent model studies suggest that GSH levels in the brain decrease with age ^4^. In contrast, human cadaver studies have shown no relationship between brain GSH levels and age in the frontal cortex and cerebellum ^5^, and GSH levels increase with age in the caudate, frontal cortex, and cerebellar cortex ^6^. Though, postmortem breakdown of GSH complicates interpretation of such findings ^7^. Magnetic resonance spectroscopy (MRS) is a noninvasive technique that can detect metabolite levels in the brain. Spectral editing techniques, such as Hadamard Encoding and Reconstruction of MEGA-Edited Spectroscopy (HERMES) ^8^, can be used to separate GSH resonance from overlapping metabolite resonances. Reproducible quantification of GSH can be achieved without the use of spectral editing techniques, such as Point RESolved Spectroscopy (PRESS) with a short echo time ^9^. However, only a few studies have examined the relationships between age and brain GSH levels in healthy participants using MRS and their findings have been inconsistent. For example, no relationship was found between age and GSH levels in the anterior cingulate cortex (ACC) in a pediatric sample (age 5.6–13.9 years) ^10^. In studies of adults, older adults showed decreased occipital GSH levels ^11^ and increased medial frontal and sensorimotor GSH levels ^12^ compared to younger adults. The regional specificity of brain GSH ^13^ and the relatively small sample sizes of prior studies are potential reasons for these inconsistent results.

In contrast, it is well established that OS increases with age and can lead to cognitive impairment ^14^, with OS playing an important role in Alzheimer’s disease (AD) onset and progression ^15,16^. Some MRS studies suggest that decreased posterior cingulate cortex (PCC) and hippocampal GSH levels are correlated with cognitive decline in patients with mild cognitive impairment ^17,18^ and AD ^19^. Notably, brain aging and AD share common clinical and neuropathological features as well as similar etiologies and pathogenesis ^20^. However, relationships between brain GSH levels, which are representative of OS, and cognitive function, have not been found in healthy participants ^11,12^. Additionally, it is known that markers of OS in the brain, such as lipoperoxidation, are higher in male brains than female brains ^21,22^. However, the effect of biological sex on the relationship between GSH levels and cognitive function has not been previously explored.

To address this research gap, this study used HERMES and PRESS to measure brain GSH levels in a large, healthy cohort and associations with cognitive function. The brain regions of interest were the ACC, which has a key role in supporting cognitive control ^23^; the PCC, which plays a significant role in spatial cognition, memory, attention, and keeping the balance between internal and external thought ^24^; and the occipital cortex (OC), which is involved in visuospatial processing ^25^. We then explored the interactive relationships between age, sex, brain GSH levels, and cognitive function. Our hypotheses included: (1) GSH levels in the brain decrease with normal aging; (2) brain GSH levels are associated with cognitive function; and (3) brain region and sex influence the relationship between GSH levels and age or cognitive function.

## Materials and Methods

### Participants

A total of 276 healthy participants (110 males and 166 females, age range 20–70 years, mean ± standard deviation = 43.63 ± 11.94 years) were recruited for this study (Table 1). All participants were of Han Chinese ethnicity, Mandarin speakers, and right-handed. The exclusion criteria were neurological or psychiatric diseases, a history of drug and alcohol abuse, contraindications for magnetic resonance imaging (MRI), and claustrophobia. This study was approved by the Shandong First Medical University Institutional Review Board. All participants provided written informed consent.

**Table 1.** Demographics.

| Variables | Overall<br>(N = 276) | Female<br>(N = 166) | Male<br>(N = 110) |
| --- | --- | --- | --- |
| <sup>a</sup>Age (years) | 43.63 (11.94) | 45.05 (11.91) | 41.48 (11.72) |
| <sup>b</sup>Education (n, %) |  |  |  |
| Illiteracy | 2 (0.7) | 2 (1.2) | 0 (0.0) |
| Primary | 18 (6.5) | 12 (7.2) | 6 (5.5) |
| Junior | 33 (12.0) | 20 (12.0) | 13 (11.8) |
| High | 39 (14.1) | 26 (15.7) | 13 (11.8) |
| AA | 41 (14.9) | 26 (15.7) | 15 (13.6) |
| College | 7 (2.5) | 5 (3.0) | 2 (1.8) |
| Bachelor | 84 (30.4) | 46 (27.7) | 38 (34.5) |
| Master | 52 (18.8) | 29 (17.5) | 23 (20.9) |
| <sup>a</sup>Cognitive function score |  |  |  |
| MoCA | 27.62 (2.66) | 27.86 (2.17) | 27.47 (2.94) |
| AVLT | 56.04 (13.60) | 56.62 (13.74) | 55.66 (13.54) |
| SDMT | 50.22 (15.17) | 51.26 (14.41) | 49.54 (15.66) |
| RCFT | 77.24 (25.83) | 80.03 (22.55) | 75.39 (27.70) |
| Stroop | 121.06 (28.73) | 121.20 (29.03) | 120.97 (28.62) |

<sup>a</sup>Mean and standard deviation (SD) are shown for continuous variables. <sup>b</sup>Frequencies and percentage are shown for categorical variables. Shapiro-Wilk test shows that age and all the cognitive function score are normally distributed.
AA, Associate of Arts; MoCA, Montreal Cognitive Assessment; AVLT, Auditory Verbal Learning Test; SDMT, Symbol Digit Modalities Test; RCFT, Rey Complex Figure Test and Recognition Trial; Stroop, Stroop Test.

### Neuropsychological assessment

All participants completed a comprehensive neuropsychological assessment taking approximately 1 h. Tests were administered in the following order: The Montreal Cognitive Assessment (MoCA) ^26^ was used to assess overall cognitive function. Auditory learning and memory were assessed using the Auditory Verbal Learning Test - Chinese version (ALVT) ^27^, attention was assessed using the Symbol Digit Modalities Test (SDMT) ^28^, working memory was assessed using the Stroop Color-Word Interference Test (Stroop) ^29^, and visuospatial memory was assessed using the Rey-Osterrieth Complex Figure Test (RCFT) ^30^.

### MR protocol

All participants were scanned using a 3.0 Tesla MR scanner (Ingenia CX, Philips) with a 32-channel head coil. Structural data were acquired using a 3D T1-weighed sequence (repetition time/echo time (TR/TE) = 8.2/3.8 ms, field-of-view (FOV) = 256 × 240 mm^2^, 160 slices with a 1 mm slice thickness, and 1 mm isotropic voxels). GSH levels were estimated using the PRESS sequence (TR/TE=2000/30 ms, bandwidth = 2000 Hz, 64 averages) and HERMES sequence (TR/TE=2000/80 ms, bandwidth = 2000 Hz, ‘‘ON/OFF’’ editing pulses = 4.56/1.9 ppm, 320 averages) in three volumes of interest (VOIs). The VOI positions are shown in Fig. 1. For the ACC, a VOI with a size of 30 × 30 × 20 mm^3^ was aligned with the corpus callosum shape and positioned superior and posterior to the genu of the corpus callosum. For the PCC, a VOI with a size of 30 × 30 × 20 mm^3^ was aligned with the corpus callosum shape and positioned posterior and superior to the splenium. For the OC, a VOI measuring 30 × 30 × 30 mm^3^ was centered within the visual cortex and aligned with the cerebellar tentorium. All VOIs were centered over the midline of the brain and placed away from the skull and sagittal sinus.

**Figure 1.**
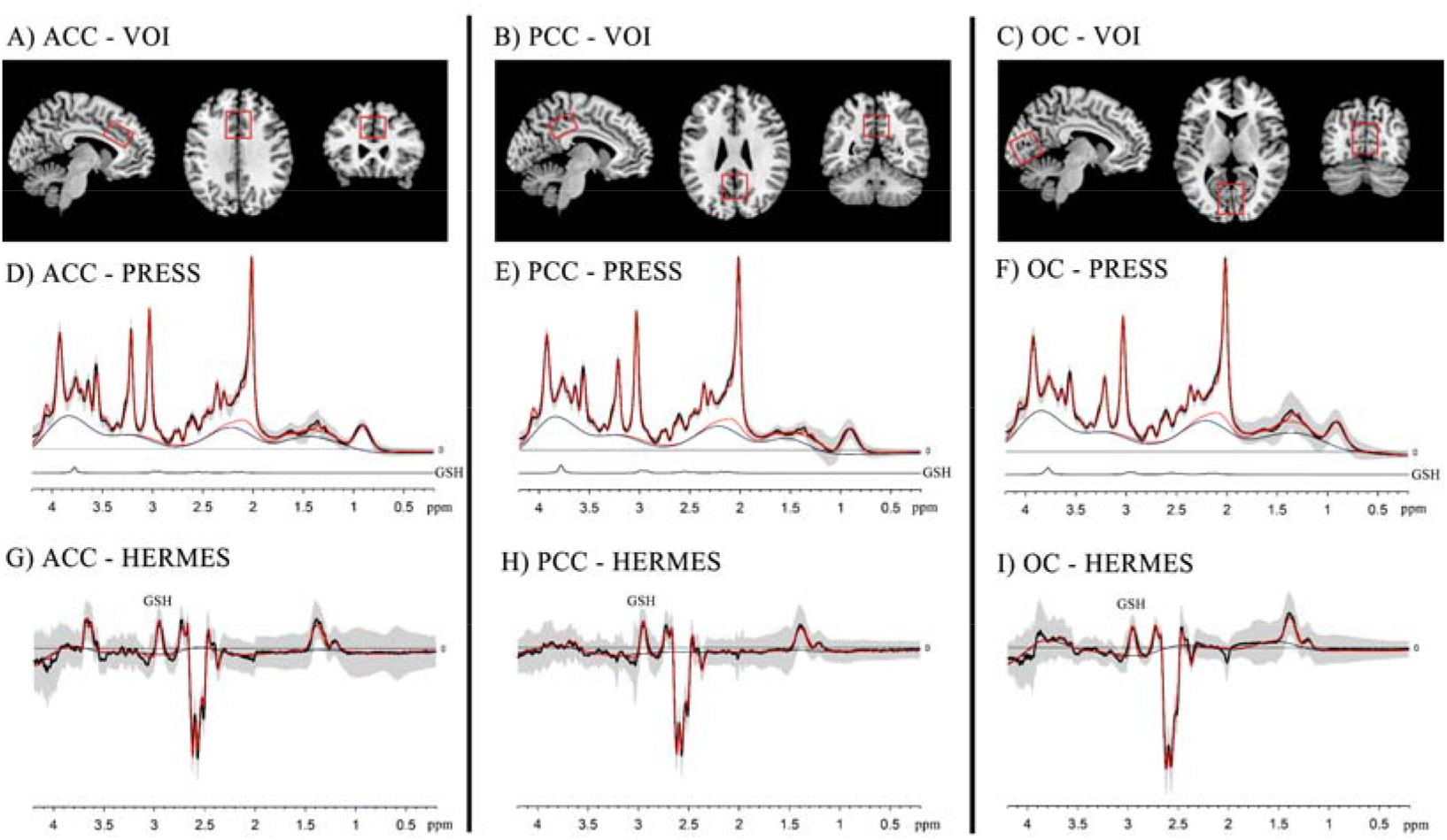
A-C) The sagittal, axial, and coronal MR images are shown, along with the VOI location, angulation, and size in the ACC, PCC, and OC. D-I) Mean processed PRESS and HERMES spectra for all individuals in the ACC, PCC and OC. Raw data is shown in black with the model fit in red and mean data ± 1 standard deviation is represented by shaded areas.

As a *J*-difference editing method, the HERMES sequence can simultaneously measure GSH and gamma-aminobutyric acid (GABA) with macromolecules and homocarnosine. Briefly, the HERMES sequence comprises four sub-experiments: A) a dual-lobe editing pulse at 1.9 ppm (ON_GABA_) and 4.56 ppm (ON_GSH_); B) a single-lobe editing pulse at 1.9 ppm (ON_GABA_); C) a single-lobe editing pulse at 4.56 ppm (ON_GSH_); and D) a single-lobe editing pulse at 7.5 ppm (OFF_GABA/GSH_). The GSH spectra were attained using the Hadamard combination A–B+C–D.

### MR data analysis

For HERMES data, Gannet 3.15 was used to analyze GSH levels with a 3 Hz line broadening ^31^. For PRESS data, GSH levels were estimated using LCModel (version 6.3-1 M) ^32^. Subsequently, individual 3D T1-weighted images were segmented using Gannet 3.15 based on SPM 12 ^33^ to derive the gray matter (GM), white matter (WM), and cerebrospinal fluid (CSF) measures of the VOIs. The partial volume effects and

T1 and T2 relaxation durations were accounted for when adjusting the metabolite levels. The water-scaled metabolite levels in mmol/kg were computed using the following equations ^34,35^:

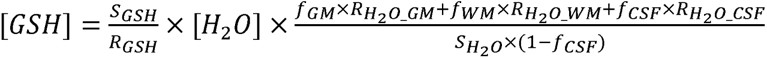

where S_GSH_ and S_H2O_ represent the GSH and water signal integrals, respectively; [H2O] is the concentration of pure water (55,550 mmol/kg); f_GM_, f_WM_, and f_CSF_ are the portions of water specifically attributed to GM, WM, and CSF, respectively ^34^; R_H2O_y_ = exp[-TE/T2_W_y_] (1 – exp[-TR/T1_W_y_]) is used to compute the relaxation attenuation factors with T1_W_y_ and T2_W_y_ being the T1 and T2 relaxation durations of water in compartment y (i.e., GM, WM, or CSF); and R_GSH_ is the relaxation attenuation factors for GSH. The following relaxation times were used: GM water: T1 = 1,331 ms, and T2 = 110 ms; WM water: T1 = 832 ms, and T2 = 79.2 ms; CSF: T1 = 3,817 ms, and T2 = 503 ms ^36–38^; GSH: T1 = 397 ms, and T2 = 117 ms ^39,40^. We excluded MRS data if the fitting errors of GSH exceeded 20%, the Cramer-Rao lower bounds of GSH exceeded 20%, or the calculated GSH levels exceeded three times the interquartile range outside the first or third quartile.

### Statistical analysis

All statistical analyses were conducted using R software (version 4.2.2) ^41^. The p-value for statistical significance was 0.05. To obtain complete data for analysis, multiple imputation was performed for excluded MRS data using the random forest method. The Shapiro-Wilk test was used to assess the normality of the variable distributions. Pearson partial correlation analysis was used to explore the relationship between age and GSH levels. To further explore potential associations, linear regression models were built. To ensure that the association effects (reflected by β coefficients) between age and GSH levels in the three brain regions measured by the two MRS sequences could be intuitively evaluated and compared, standardized age and standardized GSH levels were included in the linear regression models. Restricted cubic spline (RCS) with four knots at the 5th, 35th, 65th, and 95th percentiles were further used to identify potential nonlinearities in these associations. To remove any influence of potential confounding factors, sex, education level, full width half maximum (FWHM), and the signal-noise ratio (SNR) of total N-Acetyl-L-aspartic acid (tNAA) were included as covariates in the above analysis.

To evaluate the association between GSH levels and cognitive function scores, standardized linear regression models were built with age, education level, FWHM, and SNR of tNAA as covariates. To explore the effect of sex, the above analyses were performed separately for male and female subgroups, with age, education level, FWHM, and SNR of tNAA included as covariates.

## Results

Table 2 presents the GSH levels in the three brain regions measured by PRESS and HERMES in all participants and the male and female subgroups. Based on the criteria described in the Methods, some GSH levels measured by PRESS (ACC: 5 participants) and HERMES (ACC: 10 participants; PCC: 8 participants, and OC: 11 participants) were excluded from analysis.

**Table 2.** Distribution of glutathione in the whole population and in different sex.

| Variables | Overall<br>(N = 276) | Female<br>(N = 166) | Male<br>(N = 110) |
| --- | --- | --- | --- |
| **PRESS** |  |  |  |
| ACC | 1.93 (0.28) | 1.93 (0.28) | 1.94 (0.28) |
| PCC | 2.02 (0.29) | 2.00 (0.28) | 2.06 (0.30) |
| OC | 1.80 (0.32) | 1.79 (0.34) | 1.80 (0.28) |
| **HERMES** |  |  |  |
| ACC | 0.80 (0.34) | 0.82 (0.36) | 0.77 (0.31) |
| PCC | 1.14 (0.37) | 1.13 (0.37) | 1.14 (0.39) |
| OC | 1.14 (0.58) | 1.13 (0.57) | 1.15 (0.58) |

Mean and standard deviation (SD) are shown for continuous GSH levels.
ACC, anterior cingulate cortex; PCC, posterior cingulate cortex; OC, occipital cortex;

### Relationship between age and GSH levels

Partial correlation analysis indicates a correlation between PRESS GSH levels and age in the PCC (*r* = −0.229, *P* < 0.001), as shown in Fig. 3. Further linear regression analyses showed that PRESS GSH levels decreased with age in the PCC (β: −0.337; 95% confidence interval (CI): −0.506, −0.168; controlling for SNR, FWHM, sex, and education level). There was no significant association between PRESS GSH levels and age in the ACC (β: −0.078; 95% CI: −0.251, 0.095) and the OC (β: −0.150; 95% CI: −0.326, 0.025), as shown in Fig. 2. A non-linear relationship was not observed between age and PRESS GSH levels in ACC (*P* _nonlinear_ = 0.459), PCC (*P* _nonlinear_ = 0.466), or OC (*P* _nonlinear_ = 0.163), as shown in Fig. 4.

**Figure 2:**
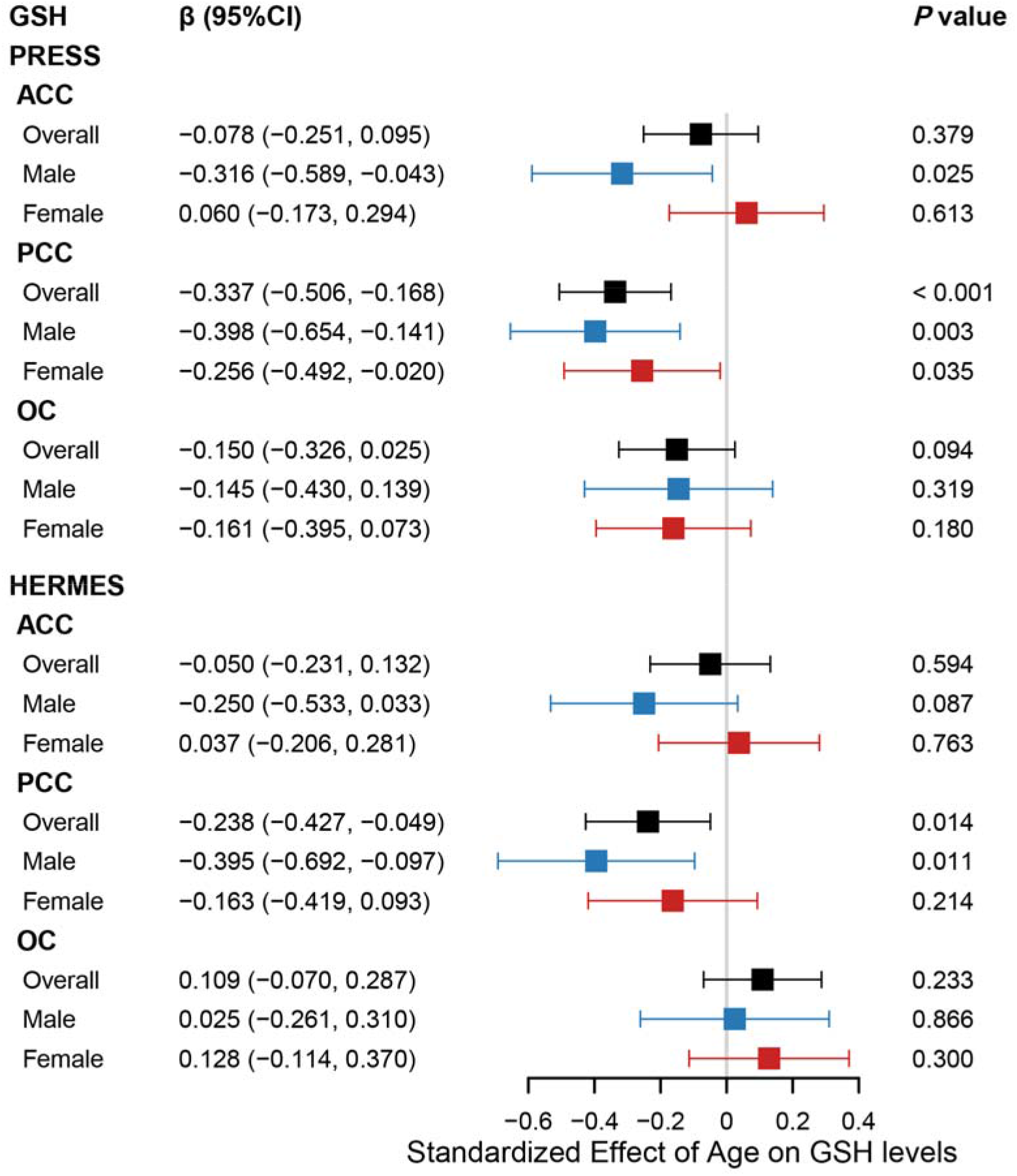
Association between age and GSH levels. Forest plot illustrated the association between age and GSH levels in overall participants, male and female. Standard linear regression model was adjusted by sex, education levels, FWHM and SNR in overall population, and adjusted by education levels, FWHM and SNR in sex subgroups.

**Figure 3:**
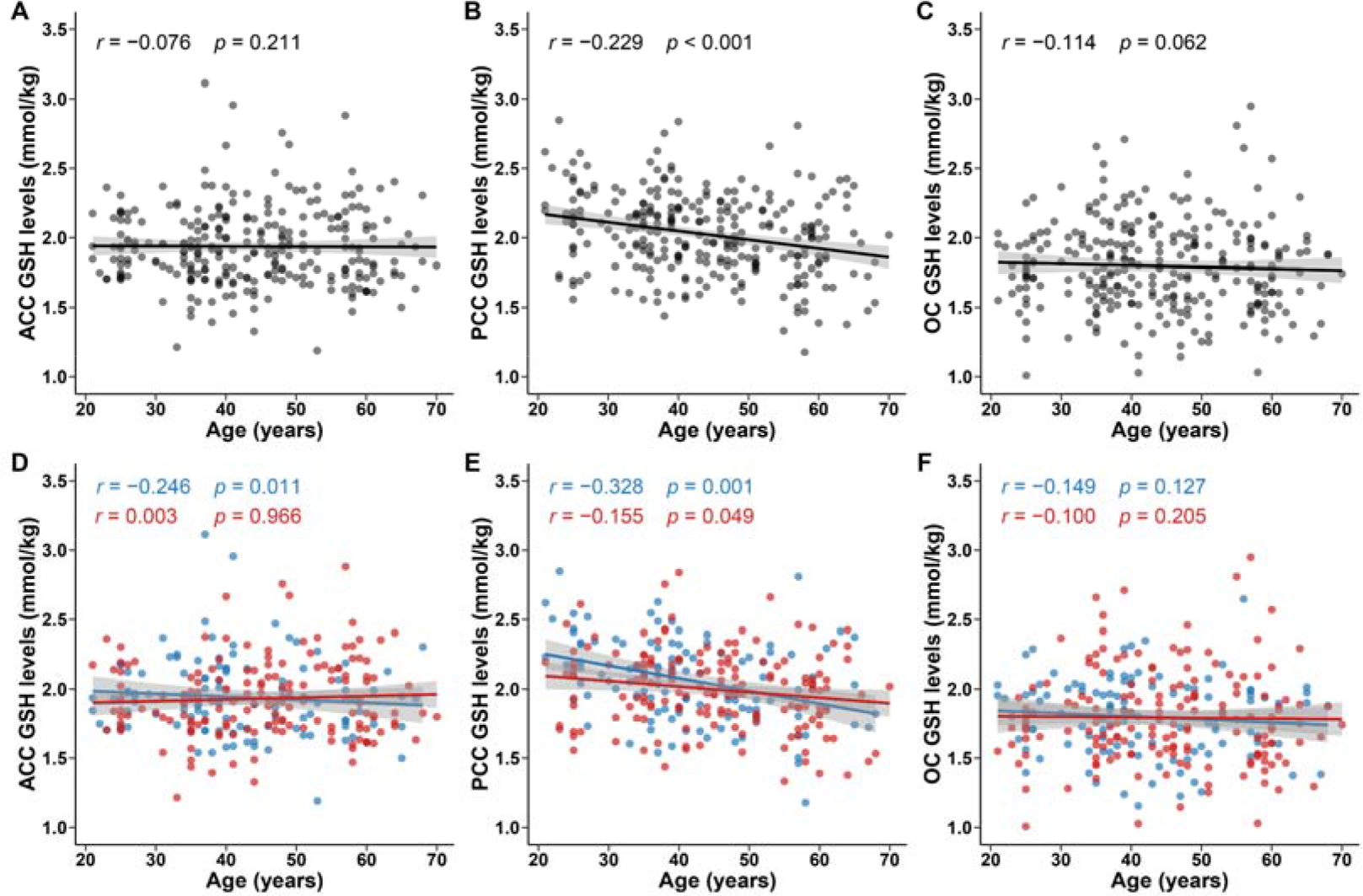
Correlation between age and GSH levels detected by PRESS. Scatter plots illustrated the distribution of age and GSH levels detected by PRESS and correlation between them. Pearson partial correlations analysis was adjusted by sex, education levels, FWHM and SNR in overall population in ACC (A), PCC (B) and OC (C), and adjusted by education levels, FWHM and SNR in sex subgroups (blue for male; red for female) in ACC (D), PCC (E) and OC (F). Fitted lines and 95% confidence intervals (shaded area) are also shown.

**Figure 4:**
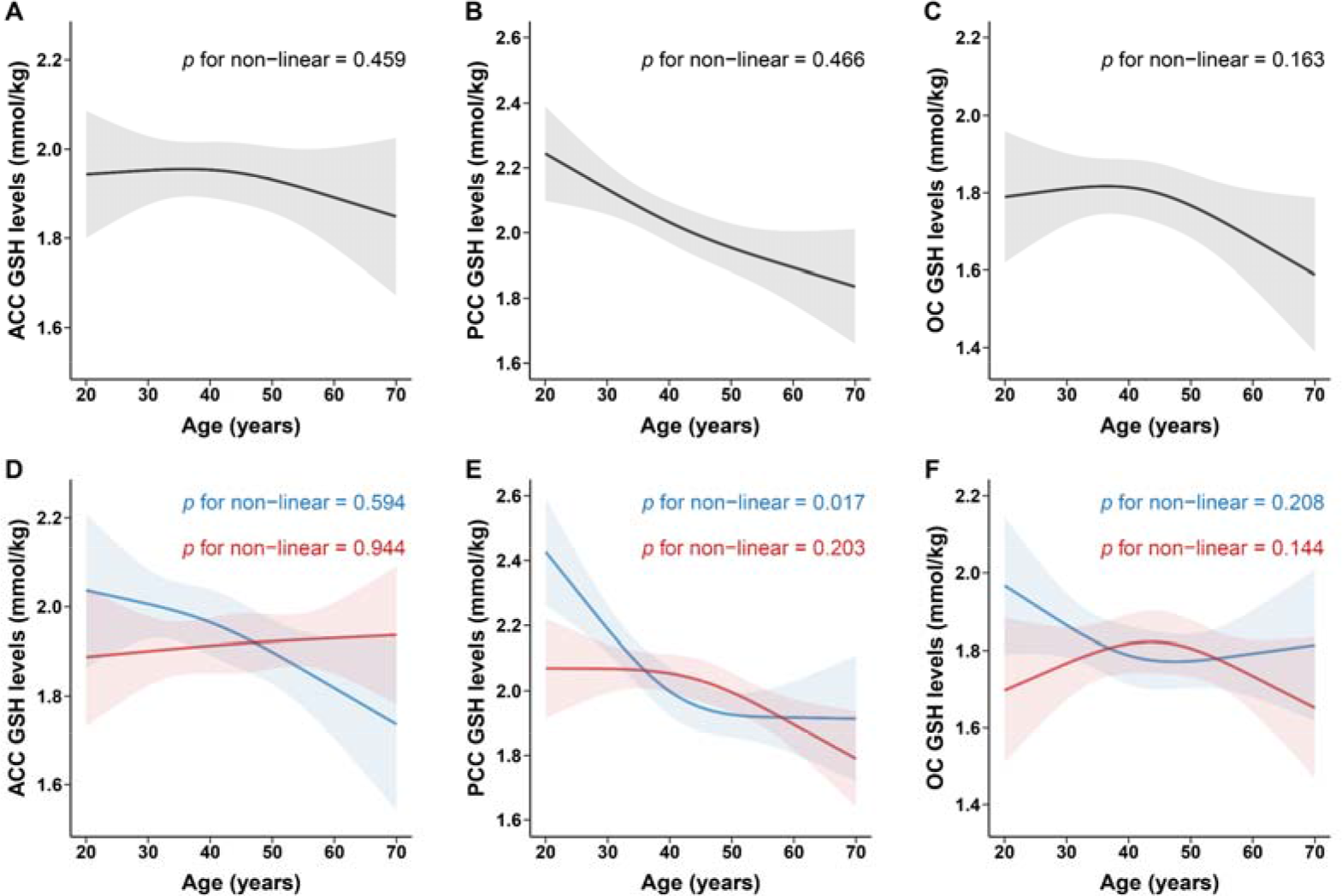
Non-linear association between age and GSH levels detected by PRESS. Restricted cubic spline was used to explore non-linear association between age and GSH levels detected by PRESS. Spline regression model was adjusted by sex, education levels, FWHM and SNR in overall population, and adjusted by education levels, FWHM and SNR in sex subgroups. All of the models with four knots located at the 5th, 35th, 65th, and 95th percentiles of age. Change of GSH levels (solid curve) and 95 % confidence intervals (shaded region) were showed in overall population in ACC (A), PCC (B) and OC (C) and in sex subgroups (blue for male; red for female) in ACC (D), PCC (E) and OC (F).

According to the edited technique, partial correlation analysis indicates a correlation between HERMES GSH levels and age in the PCC (*r* = −0.138, *P* = 0.022), as shown in Fig. 5. Further linear regression analyses showed that HERMES GSH levels decreased with age in the PCC (β: −0.238; 95% CI: −0.427, −0.049). However, there was no significant association between age and HERMES GSH levels in the ACC (β: −0.050; 95% CI: −0.231, 0.132) or OC (β: 0.109; 95% CI: −0.070, 0.287), as shown in Fig. 2. A non-linear relationship was not observed between age and HERMES GSH levels in ACC (*P* _nonlinear_ = 0.168), PCC (*P* _nonlinear_ = 0.311), or OC (*P* _nonlinear_ = 0.962), as shown in Fig. 6.

**Figure 5:**
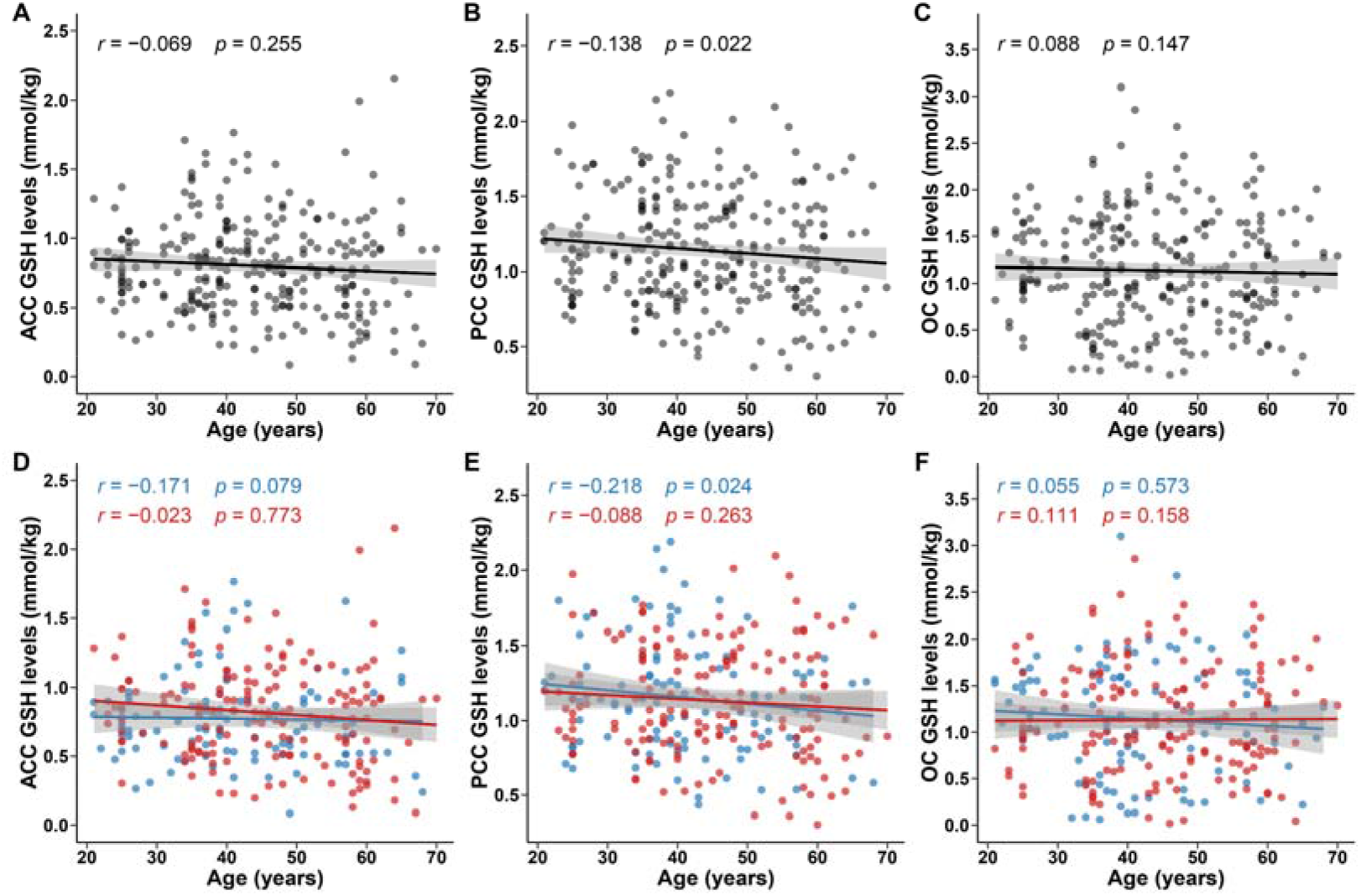
Correlation between age and GSH levels detected by HERMES. Scatter plots illustrated the distribution of age and GSH levels detected by HERMES and correlation between them. Pearson partial correlations analysis was adjusted by sex, education levels, FWHM and SNR in overall population in ACC (A), PCC (B) and OC (C), and adjusted by education levels, FWHM and SNR in sex subgroups (blue for male; red for female) in ACC (D), PCC (E) and OC (F). Fitted lines and 95% confidence intervals (shaded area) are also shown.

**Figure 6:**
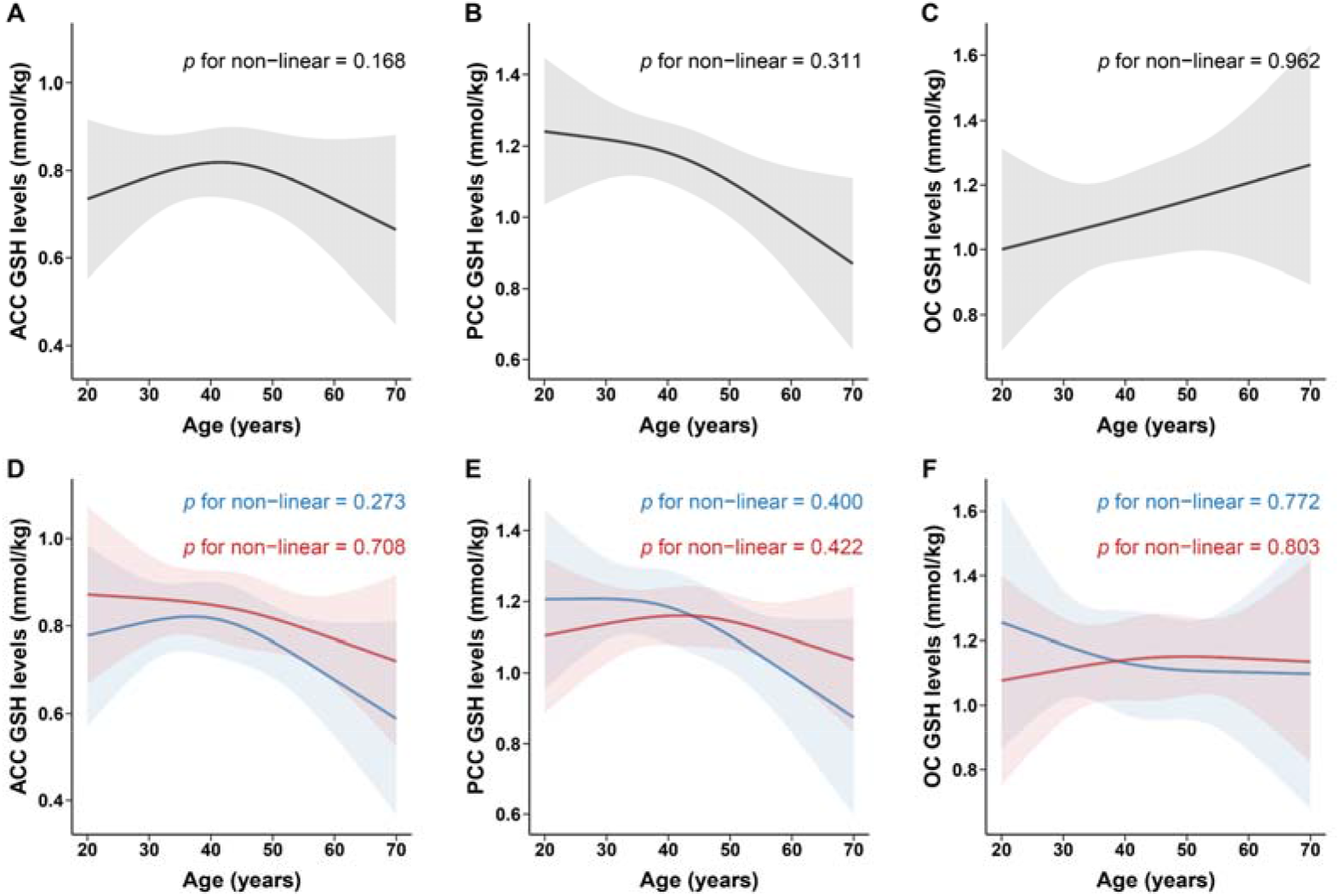
Non-linear association between age and GSH levels detected by HERMES. Restricted cubic spline was used to explore non-linear association between age and GSH levels detected by HERMES. Spline regression model was adjusted by sex, education levels, FWHM and SNR in overall population, and adjusted by education levels, FWHM and SNR in sex subgroups. All of the models with four knots located at the 5th, 35th, 65th, and 95th percentiles of age. Change of GSH levels (solid curve) and 95 % confidence intervals (shaded region) were showed in overall population in ACC (A), PCC (B) and OC (C) and in sex subgroups (blue for male; red for female) in ACC (D), PCC (E) and OC (F).

### Relationship between GSH levels and cognitive function

Linear regression analyses showed significant positive associations between RCFT scores and PRESS GSH levels in the PCC (β: 0.117; 95% CI: 0.015, 0.219) and OC (β: 0.149; 95% CI: 0.051, 0.248), controlling for SNR, FWHM, sex, and education level, but not the ACC (β: 0.102; 95% CI: 0.000, 0.203), as shown in Fig. 7. According to the edited technique, there was no significant association between RCFT scores and HERMES GSH levels in the ACC (β: −0.005; 95% CI: −0.105, 0.095), PCC (β: −0.059; 95% CI: −0.158, 0.040), or OC (β: 0.030; 95% CI: −0.070, 0.130), as shown in Fig. 7.

**Figure 7:**
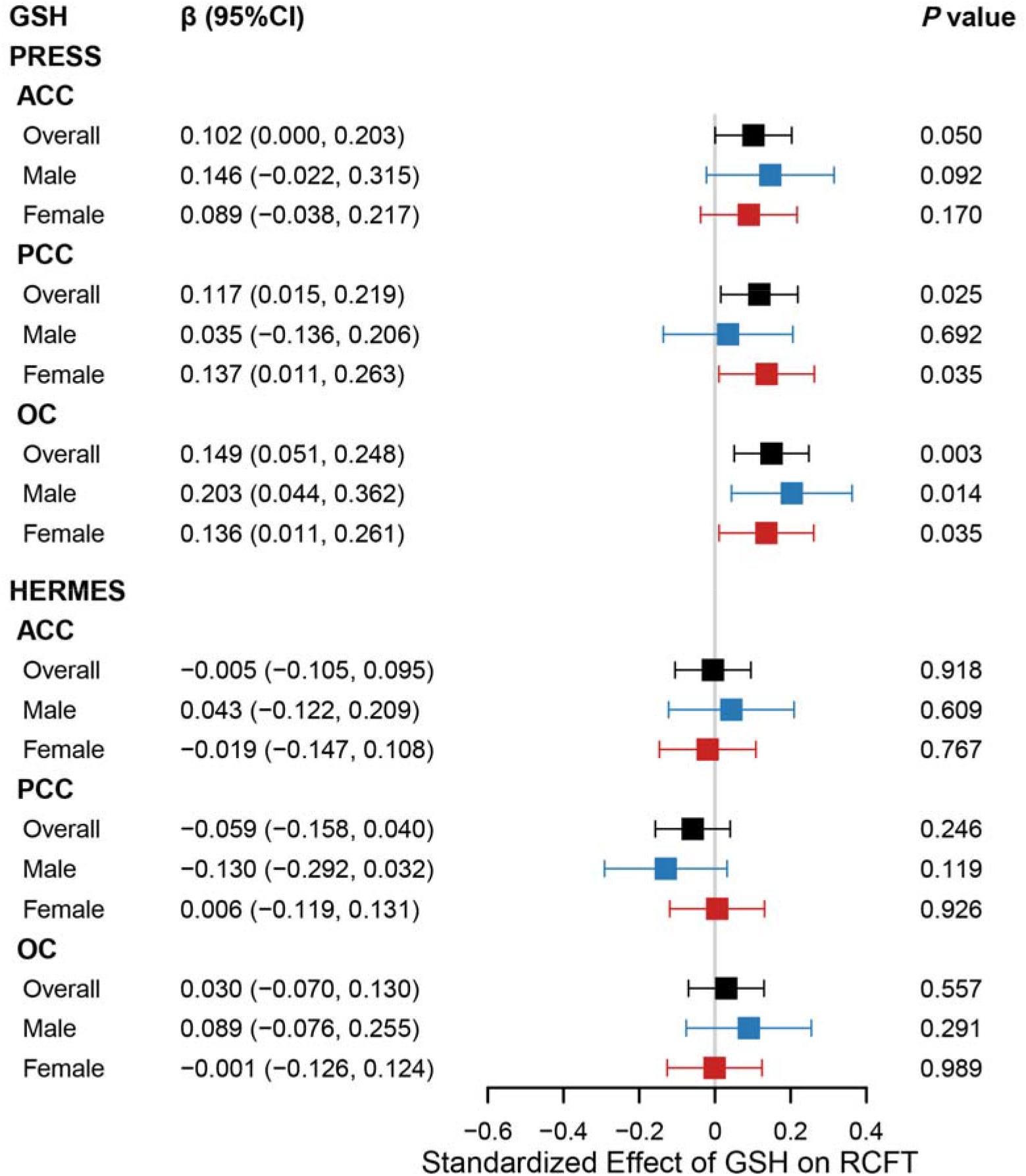
Association between GSH levels and RCFT scores. Forest plot illustrated the association between GSH levels and RCFT scores in overall population, male and female. Standard linear regression model was adjusted by age, sex, education levels, FWHM and SNR in overall population, and adjusted by age, education levels, FWHM and SNR in sex subgroups.

Furthermore, no significant associations were observed between GSH levels and MOCA, AVLT, SDMT, and Stroop scores in the total participants (see Supplementary Materials Table 1s).

### Effect of sex on the age-GSH level relationship

Further linear regression analyses of the male and female subgroups showed that PRESS GSH levels decreased with age in males (β: −0.398; 95% CI: −0.654, −0.141) and females (β: −0.256; 95% CI: −0.492, −0.020) in the PCC, as shown in Fig. 2. The correlation strength between PRESS GSH levels and age did not differ by sex (*Z* = −1.4815; *P* =0.1385). As shown in Fig. 4., a non-linear relationship was observed between PRESS GSH levels and age in males in this brain region (*P* _nonlinear_ = 0.017), but not in females (*P* _nonlinear_ = 0.203).

In the ACC, PRESS GSH levels were significantly correlated with age in males (β: −0.316; 95% CI: −0.589, −0.043), but not in females (β: 0.060; 95% CI: −0.173, 0.294). The correlation strength between PRESS GSH levels and age differed by sex (*Z* = −2.0427; *P =* 0.0411). We did not observe a non-linear relationship between PRESS GSH levels and age in this brain region in males (*P* _nonlinear_ = 0.594) or females (*P* _nonlinear_ = 0.944).

In the OC, there was no significant association between PRESS GSH levels and age in males (β: −0.145; 95% CI: −0.430, 0.139) or females (β: −0.161; 95% CI: −0.395, 0.073). We did not observe a non-linear relationship between PRESS GSH levels and age in this brain region in males (*P* _nonlinear_ = 0.208) or females (*P* _nonlinear_ = 0.144).

According to edited technique, linear regression analyses showed that HERMES GSH levels decreased with age in the PCC in males (β: −0.395; 95% CI: −0.692, −0.097), but not in females (β: −0.163; 95% CI: −0.419, 0.093), as shown in Fig. 2. The correlation strength between HERMES GSH levels and age did not differ by sex (*Z* = −1.0716; *P* =0.2839). We did not observe a non-linear relationship between HERMES GSH levels and age in this brain region in males (*P* _nonlinear_ = 0.400) or females (*P* _nonlinear_ = 0.422), as shown in Fig. 6.

No significant associations were observed between HERMES GSH levels and age in the PCC and OC in males or females and the associations were not non-linear.

### Effect of sex on the cognitive function-GSH level relationship

A positive association between PRESS GSH levels in the PCC and RCFT scores was found in females (β, 0.137; 95% CI: 0.011, 0.263), but not in males (β, 0.035; 95% CI: −0.136, 0.206), as shown in Fig. 7. The correlation strength between PRESS GSH levels and RCFT scores did not differ by sex (*Z* = −1.0802; *P =* 0.2800). RCFT scores were positively associated with PRESS GSH levels in the OC in males (β, 0.203; 95% CI: 0.044, 0.362) and females (β, 0.136; 95% CI: 0.011, 0.261), and this correlation strength did not differ by sex (*Z* = 1.0013; *P =* 0.3167).

## Discussion

To our knowledge, this is the first study to combine PRESS and HERMES to investigate age-related changes in brain antioxidant levels and their implications for cognitive function. We found that GSH levels measured by PRESS and HERMES decreased with age in the PCC. Furthermore, GSH levels measured by PRESS in both the PCC and OC were positively associated with visuospatial memory. However, only males showed a decrease in PRESS GSH levels in the ACC with age, and only females showed an association of lower PRESS GSH levels in the PCC with poorer visuospatial memory.

GSH plays a crucial role in protecting against OS and is consumed during protective antioxidant processes. In the present study, GSH levels in the PCC, measured by both PRESS and HERMES, gradually decreased with age, indicating that OS in the PCC increases with age. Consistent with our results, previous animal studies have found that aging is associated with decreased GSH levels in the rat cerebral cortex ^42,43^. As a core node of the default mode network ^44^, the PCC has the highest metabolic rate in the cerebral cortex in the resting state ^45^. As the brain ages, the PCC may become more susceptible to OS due to its high metabolism ^46^, which could result in increased GSH consumption. The close relationship between amyloid β (Aβ) and OS has been confirmed by numerous studies ^47^. Significant accumulation of fibrillar Aβ has been observed in the PCC relative to other brain regions in early-onset AD patients using positron emission tomography (PET) ^48^. Furthermore, a recent MRS-PET study found that GSH levels in the parietal lobe were negatively correlated with Aβ accumulation in healthy individuals ^49^. Taken together, these results might reflect increased consumption of GSH in the PCC during aging. In the present study, we found no correlation between GSH levels and age in the ACC and OC, which may suggest different relationships between OS and aging in different brain regions. It should be noted that there was a trend of a negative correlation between PRESS GSH levels and age in the OC, which is partially consistent with a previous MEGA-PRESS study at 4 Tesla that found decreased GSH levels in the OC in elderly healthy adults compared with young adults ^11^. In contrast, a HERMES study reported that GSH levels in the frontal lobe and sensorimotor cortex of elderly adults were higher than those in young adults ^12^. Two possible reasons for this discrepancy are the target regions of interest for spectroscopy and the sample size (37 young and 23 older adults) in the studies.

In the present study, positive correlations were found between RCFT scores and GSH levels in the PCC and OC, indicating that lower GSH levels are associated with poorer visuospatial memory. The PCC plays a critical role in spatial memory function ^50–52^. Recent studies also suggest that the PCC supports integration of visual spatial information ^53^. The OC is crucial for supporting visual spatial processing ^25^ and plays an important role in encoding visual spatial information ^54^. Previous studies have reported significant deficits in spatial memory in mice and rats after depletion of cortical GSH levels using intraperitoneal injection of 2-cyclohexene-1-one, which affects GSH levels by conjugating with the S-transferase pathway ^55^. Moreover, rescue of depleted GSH levels in the cortex by N-acetyl-L-cysteine, a GSH precursor, restored spatial memory deficits in rats ^56^. Our findings of a positive correlation between GSH levels and RCFT scores may indicate that decreased GSH levels in the PCC and OC induce OS, resulting in neuronal death and synaptic dysfunction ^14^ and ultimately leading to spatial cognitive decline. Interestingly, we found no correlations between GSH levels and MOCA or SDMT scores, which is consistent with previous MRS studies^11,12^. Thus, GSH levels in the PCC or OC could potentially serve as important predictors of visuospatial memory.

After dividing participants into two groups by sex, we found that only males showed a decrease of PRESS GSH levels in the ACC with age and a significant group difference in the relationship between GSH levels and age. This finding may reflect a weaker antioxidant defense system in the ACC in males. In line with our findings, a previous animal study reported a lower antioxidant capacity during aging in male rat brains than in female rat brains ^57^. There is evidence that the male mouse brain is more susceptible to OS due to lower levels of paraoxonase-2, which is an antioxidant enzyme positively modulated by estradiol ^58^. Moreover, paraoxonase-2 levels are substantially higher in female primate brains than in male primate brains ^59^. However, there was a significant decrease in GSH levels in the PCC with age in both males and females. As mentioned above, the PCC has a higher resting metabolic rate than other brain regions. Despite the greater antioxidant capacity of female brains, it may be difficult to balance OS due to the high-intensity metabolism of the PCC.

The present study found a significant nonlinear relationship between GSH levels in the PCC with age in males and differences in the GSH level timecourse between males and females. Specifically, in males, GSH levels decreased with age before 45 and remained stable after 45; in females, GSH levels remained stable before age 45 and decreased with age after 45. One potential cause of differences in the GSH level timecourse is the effects of sex hormones. As described above, estradiol positively regulates paraoxonase-2. During menopause, women experience a sharp decrease in estradiol secretion due to ovarian aging. Interestingly, the turning point in the GSH level timecourse of females (age 45) is within the period of menopause. Moreover, it has been demonstrated that the antioxidant defense system is impaired in female mice brains after ovariectomy ^60^. In the present study, only females showed a significant correlation between GSH levels in the PCC and RCFT scores, suggesting that female visuospatial memory is more susceptible to GSH levels. Numerous rodent studies have shown that estrogen has protective effects on spatial cognition ^61^, and estrogen is capable of reducing OS by increasing paraoxonase-2 levels. Thus, there may be a complex interaction between brain GSH levels, estrogen, and spatial cognition that warrants further investigation.

Lastly, this study is subject to several limitations. First, due to the long HERMES scanning time, GSH levels were only measured in three brain regions. As a result, several important brain regions related to cognition were not included in our study, such as the hippocampus and dorsolateral prefrontal cortex. Second, our participant sample did not cover the entire human lifespan (participants were 20–70 years old). Future studies should extend the age range or utilize meta-analysis to integrate multiple studies to investigate how brain GSH levels vary over the lifespan. Third, large volume MRS VOIs were used in this study due to the low GSH levels and resonances of GSH overlapping with other metabolites. There is potential for future MRS studies using 7 Tesla scanners to provide more detailed analyses of target brain regions by using ROIs with a smaller volume.

## Conclusion

Our findings demonstrate a decrease in GSH levels with age in the PCC, but not other brain regions. This result suggests that the PCC is more susceptible to OS during brain aging. Moreover, we found that the timecourse of GSH level changes in the PCC and ACC differed between males and females, suggesting that OS levels may be influenced by sex hormones. Importantly, positive correlations were found between RCFT scores and GSH levels in the PCC and OC after controlling for age, sex, education level, and quality of the spectroscopy, suggesting that GSH levels could serve as a predictor of visuospatial memory. Overall, these findings provide valuable insights into the relationship between brain antioxidant levels and aging or cognitive function, providing a normative foundation for investigating the neurochemical underpinnings of OS-related diseases.

## Declaration of Competing Interest

None to disclose

## CRediT authorship contribution statement

**Xin Hu:** Conceptualization, Methodology, Writing-Original Draft, Formal analysis, Data curation. **Keyu Pan:** Methodology, Formal analysis, Writing-Original Draft. **Min Zhao:** Recorded the data, Data curation. **Jiali Lv:** Software, Investigation. **Jing Wang:** Recorded the data, Data curation. **Xiaofeng Zhang:** Software, Investigation. **Yuxi Liu:** Recorded the data, Data curation. **Yulu Song:** Software, Formal analysis. **Aaron T. Gudmundson**: Edit the manuscript. **Richard A.E. Edden:** Supervision, Review, Edit the manuscript. **Fuxin Ren:** Software, Visualization. **Tao Zhang:** Conceptualization, Methodology, Supervision. **Fei Gao:** Conceptualization, Methodology, Writing - review & editing, Supervision, Funding acquisition.

## Acknowledgments

This work was supported by the National Natural Science Foundation of China (Nos. 81601479, 82222064), National Key Research and Development Program (No. 2022YFC2010100), Taishan Scholars Project (No. tsqn201812147), Shandong Provincial Natural Science Foundation of China (Nos. ZR2021MH030, ZR2021MH355), Jinan Science and Technology Development Program of China (No. 202019098), and the Academic Promotion Programme of Shandong First Medical University (No. 2019QL023). This work was also supported by NIH R01 EB016089, R01 EB023963, and P41 EB031771.

## Data Availability Statement

All data that support the findings of this study are available from the corresponding author upon reasonable request.

---

**Table 1s.** Association between GSH levels and cognitive function scores.

|  | PRESS |  | HERMES |  |
| --- | --- | --- | --- | --- |
| | $\beta^a$ (95%CI) | *P*-value | $\beta^a$ (95%CI) | *P*-value |
| ACC |  |  |  |  |
| MoCA | -0.019 (-0.119, 0.081) | 0.708 | -0.039 (-0.172, 0.094) | 0.563 |
| AVLT | -0.015 (-0.130, 0.099) | 0.793 | 0.087 (-0.066, 0.241) | 0.265 |
| SDMT | -0.004 (-0.092, 0.084) | 0.936 | -0.041 (-0.159, 0.078) | 0.502 |
| RCFT | 0.102 (0.000, 0.203) | 0.050 | 0.105 (-0.031, 0.241) | 0.131 |
| Stroop | 0.003 (-0.098, 0.105) | 0.948 | 0.045 (-0.092, 0.183) | 0.52 |
| PCC |  |  |  |  |
| MoCA | 0.076 (-0.024, 0.175) | 0.136 | -0.011 (-0.126, 0.105) | 0.856 |
| AVLT | 0.048 (-0.067, 0.163) | 0.411 | -0.013 (-0.146, 0.121) | 0.854 |
| SDMT | 0.053 (-0.036, 0.141) | 0.243 | 0.035 (-0.068, 0.137) | 0.504 |
| RCFT | 0.117 (0.015, 0.219) | 0.025 | -0.030 (-0.148, 0.089) | 0.623 |
| Stroop | -0.076 (-0.178, 0.027) | 0.150 | 0.052 (-0.067, 0.172) | 0.393 |
| OC |  |  |  |  |
| MoCA | 0.004 (-0.094, 0.102) | 0.934 | -0.045 (-0.146, 0.055) | 0.377 |
| AVLT | 0.024 (-0.089, 0.137) | 0.678 | 0.010 (-0.106, 0.126) | 0.866 |
| SDMT | 0.066 (-0.020, 0.153) | 0.135 | -0.019 (-0.109, 0.070) | 0.672 |
| RCFT | 0.149 (0.051, 0.248) | 0.003 | 0.022 (-0.080, 0.124) | 0.672 |
| Stroop | -0.014 (-0.115, 0.087) | 0.787 | 0.104 (0.001, 0.207) | 0.049 |

<sup>a</sup>Standardized coefficient of linear regression model adjusted for sex (male/female), education levels (Ordered multiple categorical variable), FWHM (continuous variable) and SNR (continuous variable). MoCA, Montreal Cognitive Assessment; AVLT, Auditory Verbal Learning Test; SDMT, Symbol Digit Modalities Test; RCFT, Rey Complex Figure Test and Recognition Trial; Stroop, Stroop Test.

## Notes

### Competing Interest Statement

The authors have declared no competing interest.

